# Characterization of a potential antibiotic target site on the ribosome

**DOI:** 10.1101/2025.05.02.651724

**Authors:** Michael Goldstein, Yang Wang, Sandra Byju, Udayan Mohanty, Paul C. Whitford

## Abstract

The ribosome is a known antibiotic target, where various classes of small molecules impact different stages of translation. There are multiple modes of function, such as interfering with the decoding process, impeding movement of the nascent protein chain and occluding the catalytic center. In the present study, we used a range of computational methods to demonstrate a new mechanism by which small molecules may impede translation in the bacterial ribosome. Specifically, we use a computational screen to identify small molecules that can bind the ribosomal protein L33 in a region that is generally conserved in bacterial species. In addition, the binding position allows it to introduce a steric obstacle that impedes the P/E hybrid-state formation steps. Using molecular dynamics simulations, we show how binding to L33 can slow down the kinetics of tRNA molecules on the ribosome. Since L33 is not present in the cytosolic human ribosome and has a distinct sequence in the human mitochondrial ribosome, this binding site may serve as a novel target for future antibiotic and antimicrobial design efforts. This analysis provides insights into how to optimize the inhibitory effects of L33-targeting molecules, in order to develop a new class of broad-spectrum antibiotics/antimicrobials.

## INTRODUCTION

The ribosome is a large biomolecular machine that is the sole producer of proteins in living organisms. Since the composition of the ribosome is organism-specific, it is also an effective target for antibiotics.^1,2^ The 70S bacterial ribosome is composed of *≈* 50 proteins and 3 RNA chains (16S, 23S, and 5S), which together form two distinct subunits (Fig. 1): the large subunit (LSU; 50S) and small subunit (SSU; 30S). During the elongation cycle, tRNA molecules and ribosome undergo a range of conformational rearrangements in order to sequentially decode frames of the mRNA and synthesize a protein.^3–6^ Aminoacyl-tRNA (aa-tRNA) is initially delivered to the A site of the ribosome by EF-Tu-GTP. Full binding of the A site requires the aa-tRNA molecule to enter the peptidyl-transferase center (PTC), a process called accommodation. After peptidyl transfer, the bound tRNA molecules are displaced from the A and P sites (Fig. 1) to the P and E sites (i.e. translocation). In order for this to occur, the P-site tRNA first adopts the so-called P/E hybrid configuration, where it moves from the P site to the E site of the LSU. Translocation is then completed when the mRNA-tRNA moves to the next binding site of the SSU. This process leads to a vacant A site, thereby allowing the next cycle of elongation to begin.

**Figure 1:**
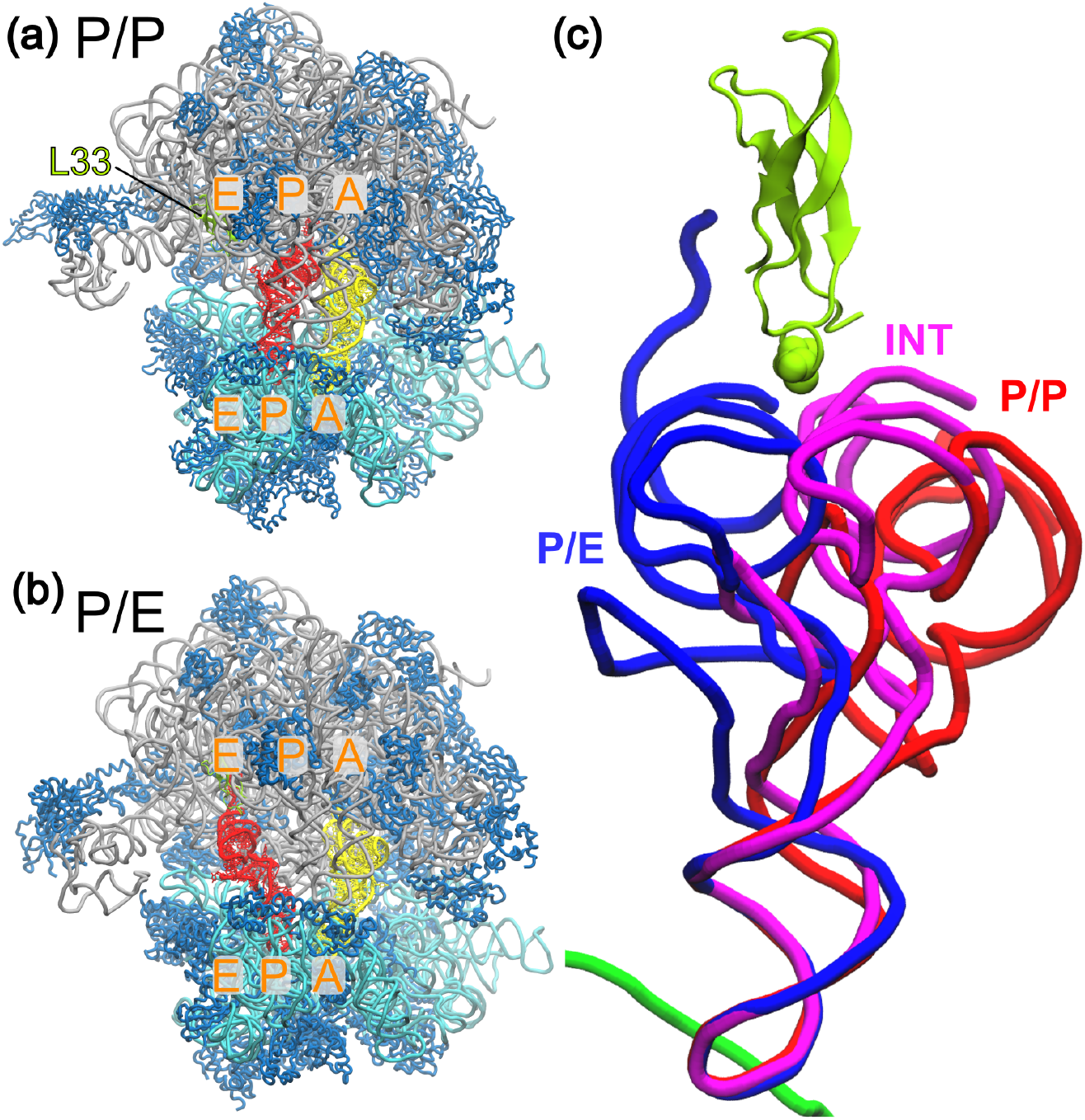
tRNA P/E hybrid state formation on the ribosome. During elongation, the ribosome undergoes a range of large-scale conformational rearrangements. After peptide bond formation, the deacylated tRNA molecule (red) must be partially displaced from the P/P conformation (panel a) and adopt the P/E hybrid state (panel b), where the tRNA molecule binds the P site of the small subunit (SSU; cyan) and E site of the large subunit (LSU; gray). c) Zoomed-in view of the tRNA molecule in the P/P (red) and P/E (blue) conformations, as well as an intermediate conformation obtained in simulations (pink). During the transition between the P/P and P/E conformations, the tRNA closely interacts with ribosomal protein L33, in particular residue R27 (shown as spheres). In the current study, we ask whether small molecules may bind near R27, in order to occlude the tRNA pathway.

Available antibiotics bind the ribosome at multiple sites and affect different stages of translation. Compounds such as edeine and pactamycin target the SSU and inhibit the initiation step,^7,8^ although their clinical use is limited due to their toxicity in model mammalian organisms. Tetracyclines, spectinomycin, and hygromycin inhibit elongation by blocking the binding of aa-tRNA to the ribosomal A site,^9–12^ while others, such as thermorubin, viomycin, and capreomycin^13–15^ interfere with tRNA-mRNA translocation. In addition, oxazolidinones, anisomycin, chloramphenicol, sparsomycin, blasticidin S, and virginiamycin M can bind the peptidyl transferase center and directly inhibit peptide bond formation.^16–18^ As a final example, erythromycin and clindamycin occlude the exit tunnel, thereby impeding movement/release of the nascent protein chain.^19,20^ While this represents a range of strategies for targeting ribosome dynamics, with the immense size of the ribosome and large number of functionally-critical conformational motions, there is significant potential for next-generation antibiotics to target novel functional sites on the ribosome.

Recent computational analysis suggested that it may be possible to target ribosomal protein L33 in order to impede the P/E hybrid state formation step.^21^ It was shown that protein L33 (specifically, the loop around residue N29 in T. Thermophilus) introduces a steric barrier during P/E formation, which suggests that binding of a small molecule could amplify the barrier. For further discussion, we will refer to this residue by its corresponding E. coli position, R27. In terms of its potential utility as an antibiotic target, the sequence of L33 is distinct between humans and bacteria, while the R27 loop region is largely conserved in bacteria. In the human cytosolic ribosome, the analogous protein is L36A, which is a circular permutant of L33 that differs substantially in sequence and structure (Figure S1). In the human mitoribosome, L33 has been found to have a similar backbone orientation as in bacteria, but the sequences are different. In contrast, the consensus sequence in bacteria is RNTPE,^21^ while in the human mitoribosome the sequence is RLREK. The consensus sequence is also very similar to that found in methicillin-resistant Staphylococcus aureus (MRSA), which is RNNPE. This relative sequence conservation in bacteria and differences with human ribosomes raise the possibility that novel broad-spectrum antibiotics may be able to bind this site in order to selectively interfere with translation in bacteria.^21^

While there has been limited attention to the biological role of L33, multiple studies indicate that it is required for proper ribosome function. When L33 is knocked out in E. coli, it presents a temperature sensitive phenotype, indicating that the protein helps ensure biological dynamics are robust to environmental changes.^22^ L33 has also been resolved in almost every experimental ribosomal structure of E. coli (SI Appendix). There is only one reported structure that lacks L33, where its absence if due to binding of the ATP-Binding Cassette F protein YbiT, which is associated with initiation (PDB 9NJV^23^). Accordingly, L33 appears to be a robust component of the ribosome that is typically present throughout the elongation cycle. Further, L33 represents a relatively rigid structural element of the ribosome, which may allow it to be amenable to small molecule binding. In terms of potential antibiotic/antimicrobial resistance mechanisms, while L33 has the potential to mutate, it is unlikely that cells can easily suppress expression. That is, L33 shares the same operon as L28, which is an essential factor for ribosome assembly.^24^ Together, these observations indicate that L33 should typically be present in the cell and associated with the ribosome.

In this study, we use a combination of molecular docking and simulation methods to demonstrate the potential of L33 as an antibiotic/antimicrobial target. In this multi-step approach (See Fig. S2), we first performed a computational screen for molecules that are predicted to bind L33. From this, we identified several small molecules that are likely to bind L33 in positions that would allow them to interfere with hybrid formation dynamics. We additionally simulated P/E formation in the presence of the bound molecule, which shows that it can significantly reduce the rate of P/E formation. This reveals specific molecular properties that may be leveraged to design effective antibiotics that bind tightly and have strong inhibitory effects. Since no drugs are known to block translation by inhibiting P/E hybrid formation, the identification and targeting of this site may enable development of a new class of antibiotics and/or antimicrobial agents.

## RESULTS

### Computational screen identifies molecules that can bind protein L33

In order to identify possible strategies for targeting elongation dynamics, we first performed a computational screen to identify small molecules that are likely to bind ribosomal protein L33. Since L33 interacts closely with the tRNA molecule during P/E hybrid formation, we focused on the solvent exposed surface, where binding could introduce a steric barrier to movement of tRNA (Fig. 1). For this, we performed molecular docking calculations for *≈* 3.5 *×* 10^8^ drug-like molecules that are predominately neutral at pH 7.4.^25^ (see methods). Neutral molecules were selected because negative charges can reduce membrane permeability, and positively charged molecules are unlikely to bind the positively charged L33 protein, estimated to be +7 in neutral aqueous solution by the ff14SB forcefield. ^26^ For docking calculations, the structure of L33 was obtained from a structure of the complete E. coli ribosome (RCSB ID: 4V9D^27^). To identify molecules that can bind the region of interest, calculations were initially performed using only the region around R27 (Fig. S3), which was previously predicted to contact tRNA during hybrid formation.^21,28^

The docking calculations resulted in a wide range of scores that spanned from unfavorable (e.g. highest score) to favorable values (−57; Figure 2). We excluded the best scoring molecule because the estimated partial charge assigned in the ZINC database includes a highly positive thioether sulfur (+3.8) that is likely to be unphysical. Of the remaining molecules, we considered those with a score of -33 or better for further analysis. A cursory glance reveals some common molecular properties. Specifically, the molecular weights of these molecules ranged from 350 to 500g/mol, while the LogP (measurement of “greasiness”) ranged from -1 to 3.5. The relatively low LogP values (all well below 5) indicate potential for future synthetic modification to improve affinity or pharmacokinetic properties while remaining feasible as potential drug molecules.^29^ As described below, this set of molecules (*N* = 24, 235) was then subjected to additional tests to determine whether they are likely to block tRNA motion during P/E hybrid formation.

**Figure 2:**
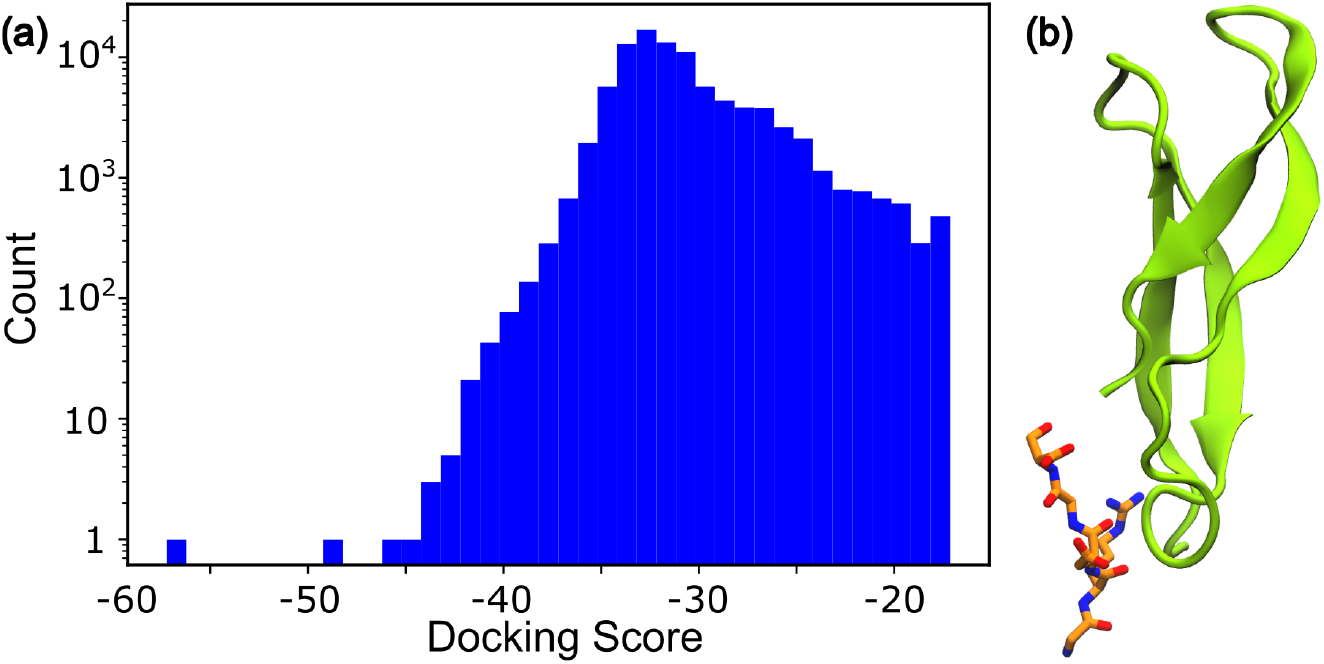
Small molecules predicted to bind L33. a) Distribution of predicted binding scores for the 1000 top-scoring molecules, out of *≈*3.5 *×* 10^8^ molecules in the ZINC database.^25^ b) Many molecules are predicted to have an affinity for L33 (green) in the vicinity of R27. As an example, the molecule shown had a score of -48.

**Figure 3:**
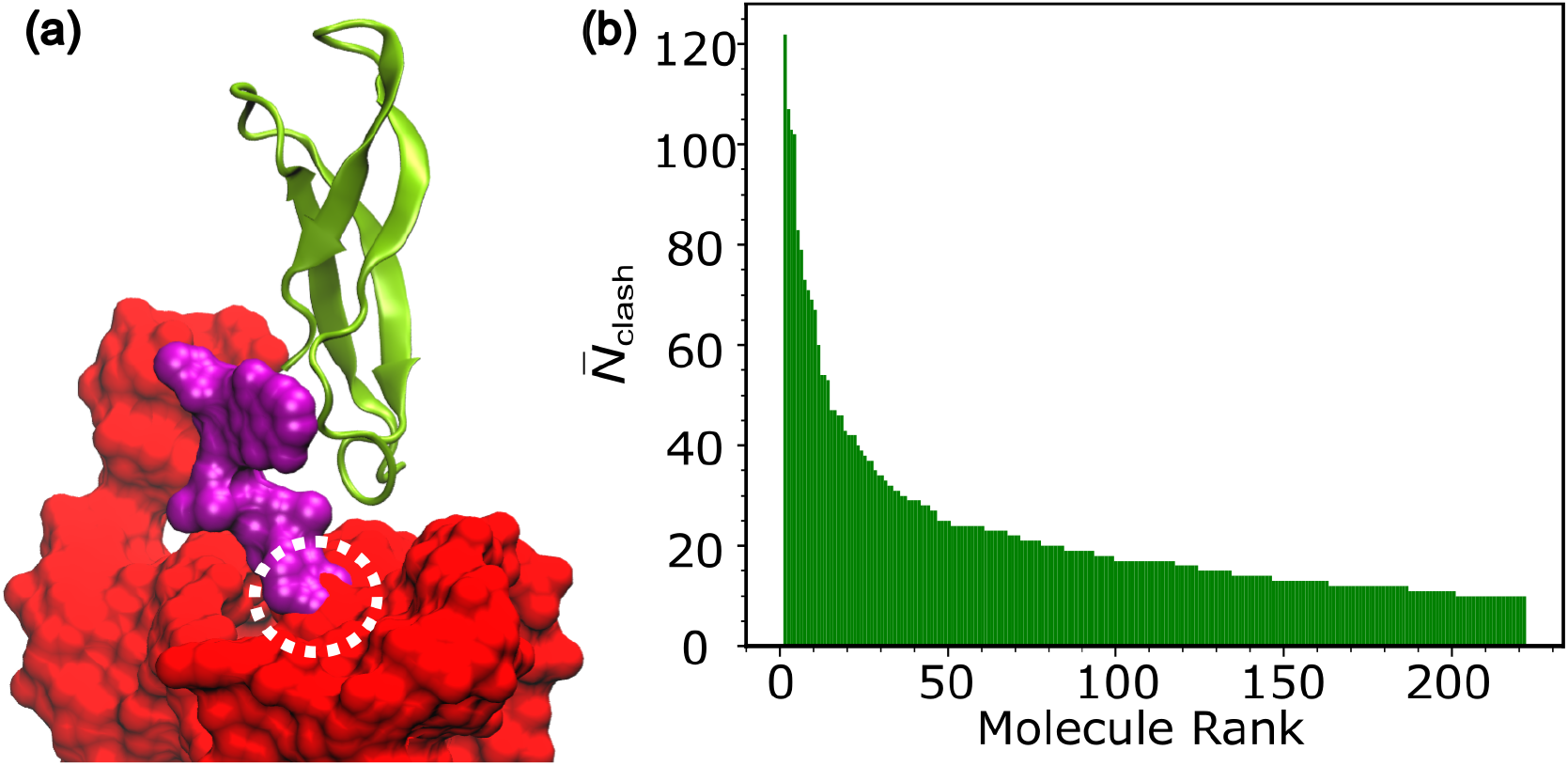
Binding poses overlap with path of tRNA during P/E formation. a) Representative small-molecule binding pose, overlayed with a snapshot obtained from a simulation of P/E formation. There is distinct overlap between the molecule and tRNA, suggesting it may be able to impede P/E hybrid formation on the ribosome. b) Average number of atomic clashes 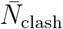, calculated for each small molecule and averaged over 45 simulations of P/E formation. 222 molecules were associated with 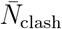 values greater than 10.

**Figure 4:**
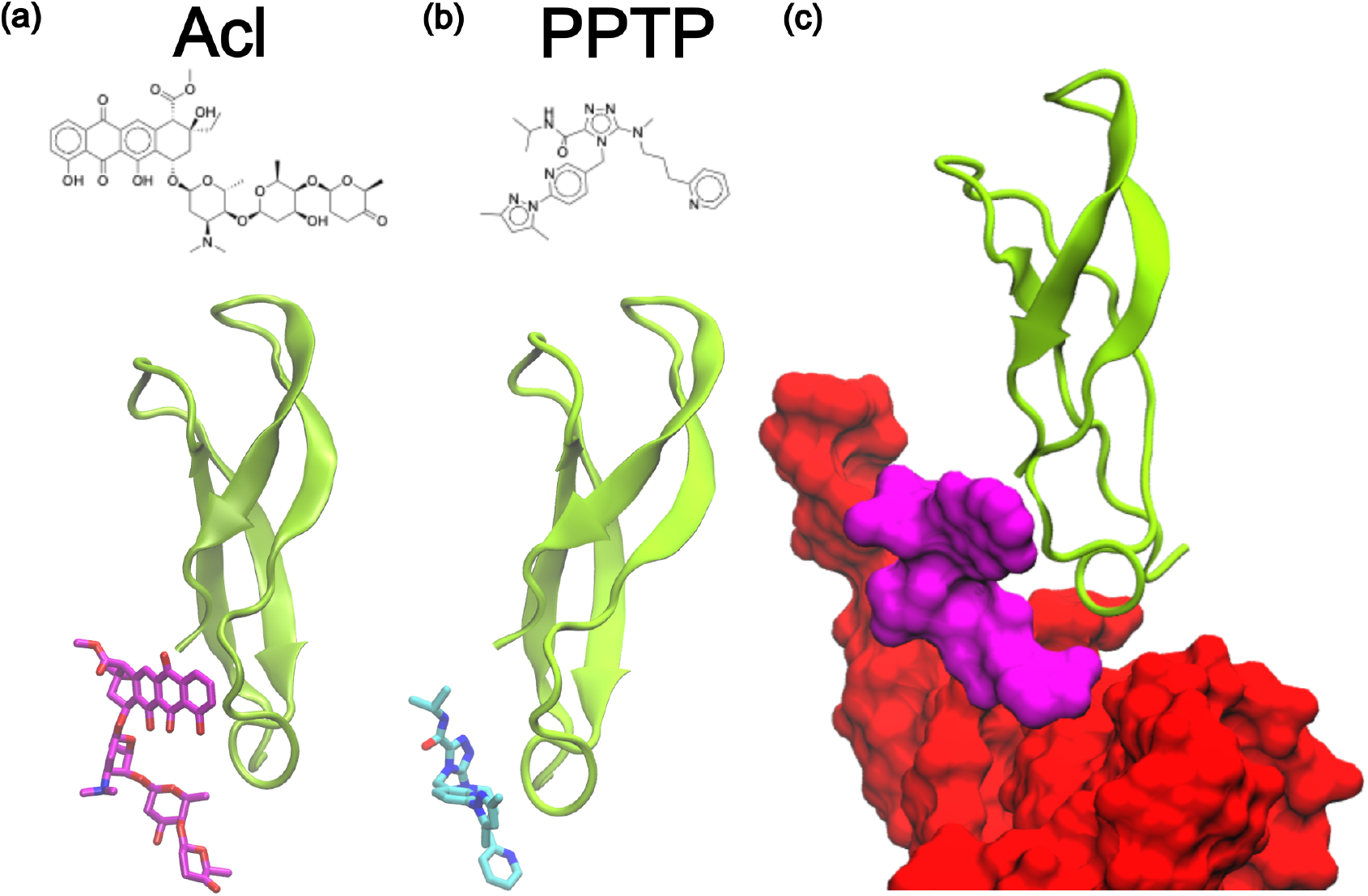
Top candidate molecules for P/E hybrid formation inhibition. For molecules that yielded 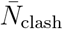 values greater than 10 (Fig. 3), docking calculations were repeated using an expanded region of L33 (Fig. S3). 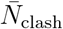 values were then recalculated for the new binding poses. After redocking, two molecules were found to bind in positions compatible with P/E inhibition 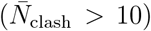: Aclurubicin (Acl; panel a) and a pyrazole pyridine tetrazole pyridine molecule (PPTP; panel b). Simulations were then performed with Acl or PPTP bound to L33. c) Representative simulated snapshot that shows Acl is positioned in a manner sufficient to impede tRNA motion during P/E formation.

### Small molecules bind along the tRNA pathway in the ribosome

For molecules that were predicted to bind L33, we next asked if their binding poses could introduce a steric obstacle during P/E tRNA hybrid formation. For this, we used an allatom structure-based (SMOG) model^30^ to simulate movement of tRNA molecules between the P/P and P/E conformations. As described previously,^21,28^ this model is constructed to have two dominant potential energy minima that correspond to the A/A-P/P and P/P-E/E conformations. In this model, experimental structures are used to define the endpoints. While the P/E hybrid configuration is not explicitly defined to be a minimum, the combined interactions with the P site of the SSU and E site of LSU allow this state to also be stable. With this simplified energetic scheme, the primary objective is to identify sterically-induced barriers that arise from molecular structure. In previous applications, these models were able to identify a precise steric contribution of protein L33 during hybrid formation, where similar models have been able to identify sterically-induced barriers during aa-tRNA accommodation^31,32^ and tRNA-mRNA translocation.^33^ Accordingly, by using a structure-based model, we specifically tested whether the binding of a small molecule can introduce a steric hindrance that will impede P/E hybrid formation.

Before discussing how specific molecules may interfere with P/E formation, it is worthwhile to briefly describe the overall character of the simulated transitions. We monitored P/E hybrid formation using three reaction coordinates: *R*_P_, *R*_E_, and *ρ*_1,2_ (See methods and Fig. S4). The dynamics with this model were consistent with a similar model that was applied to study P/E formation in a T. Thermophilus ribosome. ^21^ That is, hybrid formation occurs via similar intermediates, which are referred to as I1 and I2. (Figure S5) The I1 intermediate is associated with initial dissociation of tRNA from the P site of the LSU. To reach the I2 ensemble, the tRNA molecule must transition past protein L33 and rRNA helices H74/H88. (Figure S5) It was previously shown that the rate-limiting step in these simulations is the transition between the I1 and I2 states. Accordingly, our analysis focuses on molecules that may bind L33 and amplify this rate-limiting barrier.

Using our simulations of P/E formation, we determined whether each predicted binding pose would lead to steric interactions between the small molecule and tRNA. To this end, we first aligned the predicted ligand-L33 complex to every frame of a simulated trajectory, where alignment was performed based on the *C*_*α*_ atoms. We then counted the number of times that the aligned small molecule position was within 0.5 Å (i.e. a “clash”) of the tRNA molecule when transitioning to the P/E configuration. As a proxy measure for the ability of a molecule to impede P/E formation, we calculated the average number of atomic clashes that were identified in a simulation, 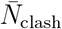. This analysis was performed separately for every small molecule that had a relatively low binding score (identified above).

For molecules that had a large number of clashes (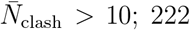molecules), we repeated the docking calculations using an expanded region of L33 (Figure S3). This additional docking step was performed to determine whether the blocking-capable pose of each molecule was also the favored position. Unfortunately, most molecules heavily favored alternate poses/regions. However, we found that three molecules have dominant poses in the site of interest. For these, we again repeated the clash-based analysis and found that two have positions that allow for interactions with the tRNA molecule. Specifically, Aclurubicin (Acl) and PPTP (Pyrazole Pyridine Tetrazole Pyridine; ZINC001304731947) have 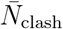 values of 40.2 and 11.7. Interestingly, these molecules have similar LogP values of 3.16 (Acl) and 3.13 (PPTP), while the molecular weights are 812 and 488. While we do not expect these specific molecules to serve as antibiotics (see discussion below), this analysis does suggest a general scale and classes of molecules that may be more effective at halting P/E formation.

### Small molecule binding can impede tRNA kinetics

As a final test of the identified drug-like molecules, we performed simulations with the either Acl or PPTP bound to L33. For this, we defined the predicted binding poses as stable, so that the small molecules remain in their predicted binding positions. We then simulated the ribosome with each small molecule bound and compared the mean first-passage time for P/E formation with a ligand present or absent (Fig. S6). While PPTP had a negligible effect on P/E formation kinetics, Acl slows the kinetics by a factor of *∼*3 (Figure 5). Accordingly, this analysis shows that Acl-like molecules may be capable of targeting the bacterial ribosome by specifically interfering with the P/E hybrid state formation step.

**Figure 5:**
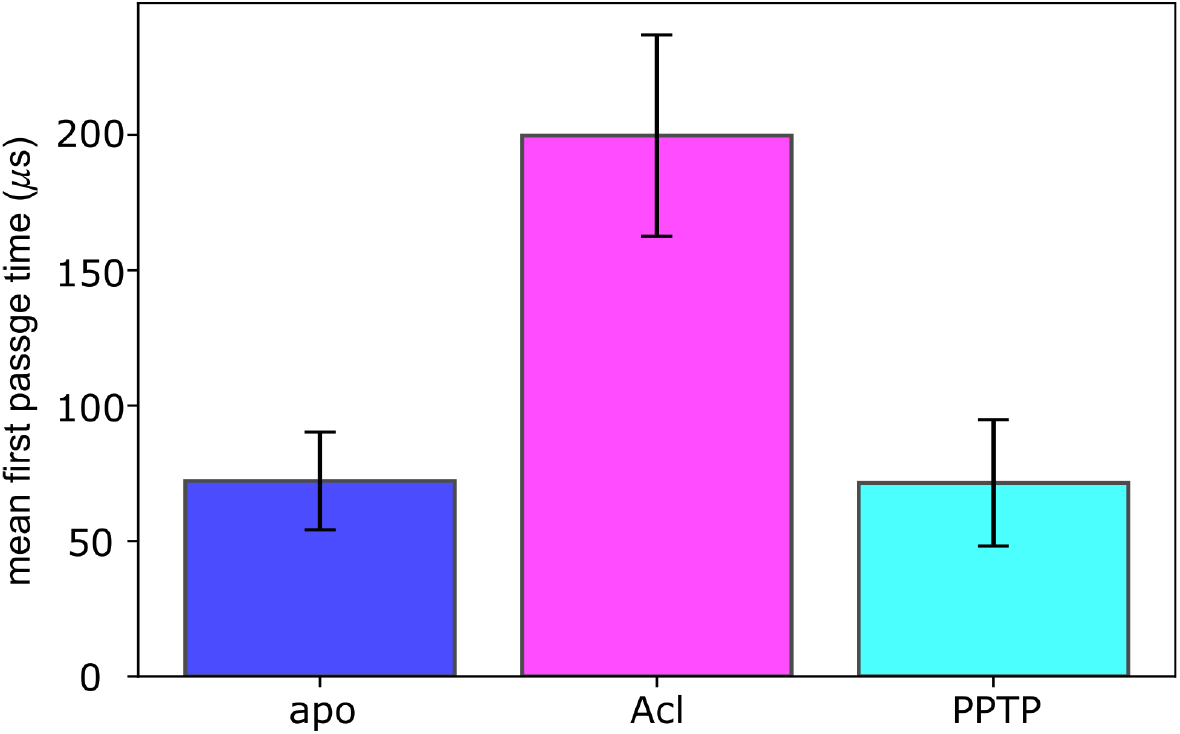
Small molecules can slow down P/E formation kinetics. To test whether Acl or PPTP is able to impede P/E hybrid formation, the mean first-passage time for P/E formation was calculated in the absence (apo) and presence of the small molecules. Acl is found to significantly slow down P/E formation. This suggests that Acl-like molecules may be designed and optimized for P/E inhibition. Error bars represent standard error, calculated from multiple simulations of each system.

Intriguingly, Acl is an anthracycline anticancer drug that is known to intercalate with DNA and block the action of topoisomerase II, where the aromatic portion of the molecule intercalates and the glycan tail allows it to further disrupt the DNA double helix structure.^34,35^ It also has antimicrobial activity through the same mechanism. While experimental validation is needed to confirm binding of Acl to protein L33, the presented analysis suggests some general considerations that may be used to design new antibiotics that target the P/E formation step. Like Acl, a ligand with an inflexible aromatic “head” can fit into the slight groove formed by K26, P30, E31, K32 and K35. In addition, a more flexible tail can wrap around and contact R27, allowing it to remain bound to the L33 protein in a manner that occludes the path of the tRNA (4). Consistent with the clash-based analysis above, these full-ribosome simulations demonstrate the ability of a small molecule to protrude from the L33 protein and impede P/E formation.

To demonstrate how one may be able to use Acl-like molecules to design new ribosome-targeting antibiotics, we also considered a hypothetical molecule. Since the current study seeks to identify molecules that can sterically occlude P/E formation, we considered the impact of adding a bulky group to Acl. Specifically, we constructed a model of Acl that has a bulky cyclohexyl group attached to its glycan tail (Figure S7). In this hypothetical molecule, the ketone oxygen is replaced by an ether oxygen, to which the cyclohexyl group is attached. We then considered the predicted binding pose of Acl and constructed an energetic model in which this composite molecule binds L33, while carrying a larger excluded volume. In this model, all other simulation parameters were unaltered. Consistent with the results obtained for Acl, the addition of the cyclohexyl group further increased the timescale for P/E hybrid formation, where it was nearly 4 times longer than the ligand-free simulations.

## DISCUSSION

A major goal in the study of ribosome dynamics has been to identify pathogen-specific molecular properties that may be exploited to develop novel therapies that have minimal side effects. While biochemical, single-molecule and structural studies have provided a wealth of information about the details of ribosome function, these methods primarily describe the properties of longer-lived states. To complement such insights, theoretical models may now be used to isolate transient interactions that are formed during large-scale conformational motions.^36^ With this extended view of ribosome dynamics, there are new opportunities to target stages of function in a precise manner, which provides a foundation for the design of next-generation antibiotics and antimicrobials.

In the current study, we show how insights into the dynamics of tRNA motion on the ribosome may be leveraged to identify small molecules that can interfere with ribosome dynamics in bacteria. Specifically, we provide evidence that an available small molecules (Acl) may bind ribosomal protein L33 in a position that allows it to impede movement of tRNA molecules. With this proof-of-principle example, there are many additional directions that may be applied to develop effective therapeutics. As an example, one may aim to design Acl derivatives that have increased binding affinity, stronger inhibitory effects and minimal side effects.

Unlike many ribosome-binding antibiotics, the target site described here is composed entirely of amino acids. Accordingly, this may represent a target that is also amenable to binding of antimicrobial peptides. While we used a screen-based approach to identify small molecules that can target L33, we anticipate that recent advances in machine learning may allow for efficient design of small peptides that exhibit antimicrobial properties. Further, while L33 is relatively conserved across known bacteria, there may also be opportunities to design pathogen-specific peptides that are tuned to subtle variations in its sequence. In future studies, it will be exciting to see how a combination of theory, computation and experimental techniques may be leveraged to devise new strategies for combating evasive bacterial species.

## MATERIALS AND METHODS

### Small molecule docking

All docking calculations used a structure of L33 protein found in a crystal structure of the E. coli ribosome (PDB ID 4V9D^27^). Docking calculations were performed using Dock6.^37^ For this, molecular surface and respective normal vectors of the protein were calculated using a probe radius of 1.4 Å. Surface spheres were then computed by aggregating surface points to atoms on the solvent accessible surface, with 1.4 Å *≤* radius *≤* 4.0 Å. Spheres around the region of interest (within 8.0 Å of residue ARG27) were then selected for the first pass at docking. A box was then constructed around the region of interest, only encompassing the solvent accessible face of the protein. The dimensions of the initial box were 26 Å x 22 Å x 22 Å and the extended box had dimensions of 22 Å x 23 Å x 48 Å (Fig. S3). For all docking calculations, the non-bonded energies were discretized on a grid with spacing of 0.14 Å. For protein electrostatics and vdW forces, AMBER99 parameters^38^ were used. 3D structures of ligands were downloaded from the ZINC database.^25^ Molecules were restricted by the “drug-like” categorization, which relies upon Lipinski’s rule of 5,^29^ as well as an added restriction of overall neutral charge at pH 7.4. This included *≈* 3.5 *×* 10^8^ molecules, which were all docked against L33. For the ligand orientation, the anchor-and-grow algorithm was used.

### Simulation of P/E hybrid formation

All simulations used an all-atom structure-based force field.^30^ The potential energy of the tRNA was defined to have two dominant minima, corresponding to the A/A-P/P (pre) and P/P-E/E (post) configurations.^28^ As described in the main text, this representation leads to a stable P/E conformation, where stabilizing interactions are simultaneously formed with the P site of the SSU and the E site of the LSU. Force field files were generated with SMOG 2.4.^30^ Simulations were performed using the OpenSMOG^39^ libraries with OpenMM.^40^ Langevin dynamics were used to maintain a temperature of 0.5 reduced units and an implicit solvent. Analysis of trajectories using the MDtraj library.^41^

The multi-basin model is constructed such that the P-site tRNA has stabilizing contacts between both the P-site and E-site on the large subunit, which allows the tRNA to undergo spontaneous (i.e. non-targeted) hybrid formation events. Complete model description provided in Ref.^28^. Every simulation was initiated from the A/A-P/P structure. All intra-ribosome contacts and dihedrals are assigned based on the reference rotated structure (PDB 4V6E^42^). All tRNA and mRNA contacts and dihedrals are based on the A/A-P/P configuration. The P/E state of the tRNA is stabilized by the addition of contacts between E-tRNA and E site based on the POST translocation P/P-E/E structure. Since SBM describes effective energetics, contacts that are formed transiently are assigned weaker strength. The contacts formed by the P-tRNA with the LSU are assigned a weaker strength compared to the intra-ribosome contacts. The contacts between the P-tRNA with the P site from the A/A-P/P configuration are assigned a contact strength of 0.05 *ϵ*_C_, where *ϵ*_C_ is the default strength of intra-ribosome contacts. To ensure stable binding of the tRNA with the E site during hybrid formation, the contact strength between A76 of CCA tail and residues G2421/C2422 of the 23S rRNA (LSU) is assigned a stronger contact strength of *ϵ*_C_. All other contacts between the tRNA and E site were given a weight of 0.25 *ϵ*_C_. Harmonic restraints (spring constant of 2 reduced units) were introduced between a base pair contacts in the P-tRNA acceptor arm, specifically between residues G1 and C72. This was added to prevent artificial unfolding of the tRNA acceptor region due to low contact density in this region. Harmonic potentials were assigned for bonds, bond angles and rigid dihedrals (e.g. rings). Flexible dihedral angles were represented by cosine functions. Atoms in contact with one another, as defined by the Shadow algorithm,^43^ were given attractive 6-12 Lennard-Jones-like potentials. All other non-bonded interactions were given a repulsive excluded volume term, to ensure a proper treatment of molecular sterics. P/E hybrid formation events were identified based on three reaction coordinates: *R*_P_, *R*_E_, and *ρ*_1,2_ (Fig. S4).

The same force field was used to compare the ligand-free and ligand-bound dynamics. In the ligand-bound simulations, the bound pose of the small molecule was defined to be the lowest-energy configuration, where contacts between the small molecule and L33 ensured that the binding pose was maintained.

For all simulations, the reduced time unit was estimated as corresponding to 1 nanosecond, as estimated previously for SMOG models applied to tRNA inside of the ribosome.^44^

### Calculating 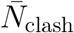

Simulations of the ligand-free system were used to determine whether a predicted binding pose could introduce a steric obtacle during P/E formation. For this, L33 and P-tRNA coordinates were extracted from 45 different P/E hybrid forming trajectories. Ligand-docked L33 structures were aligned to the L33 conformation in each trajectory. The distances between all ligand atoms and the tRNA were then calculated for each simulated frame. If an atom-atom distance was below 0.5 Å, it was marked as a “clash,” indicating that the ligand would interact with the tRNA and possibly impede P/E hybrid formation. 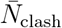 the number of clashes per simulation, averaged over all 45 trajectories.

### Enumerating available L33 structures

A list of all known E. coli ribosome structures was enumerated using the Ribosome Analysis Database (http://radtool.org,^45^). This list was then used to query in the RCSB database. In-house scripts were used to identify which structures contained the L33 protein. The complete report is provided in Appendix A.

## ACKNOWLEDGMENTS

This work was supported by the National Institutes of Health and the National Science Foundation.

## DATA AVAILABILITY STATEMENT

Simulation data and analysis scripts will be made available, upon request.

