## Supplementary Information for "Characterization of a potential antibiotic target site on the ribosome"

### Supplementary Information for:
### Characterization of a potential antibiotic target site on the ribosome

Michael Goldstein[1], Yang Wang[1], Sandra Byju[2,3], Udayan Mohanty[1], and Paul C. Whitford[2,3]

[1]*Department of Chemistry, Boston College, Merkert Chemistry Center, 2609 Beacon St, Chestnut Hill, MA 02467, USA*
[2]*Center for Theoretical Biological Physics, Northeastern University, 360 Huntington Ave, Boston, MA 02115, USA*
[3]*Department of Physics, Northeastern University, Dana Research Center 111, 360 Huntington Ave, Boston, MA 02115, USA*

#### List of Figures

S1 **Bacterial L33 and equivalent proteins in human ribosomes.** . . . . . S2
S2 **Summary of the workflow for this study.** . . . . . S3
S3 **Boxes used for molecule docking in this work.** . . . . . S4
S4 **Reaction coordinates used to detect P/E hybrid formation.** . . . . . S5
S5 **Intermediates seen along the route to P/E hybrid formation.** . . . . . S6
S6 **Representative time traces of SBM trajectories.** . . . . . S7
S7 **Acl with a cyclohexyl group attached.** . . . . . S8

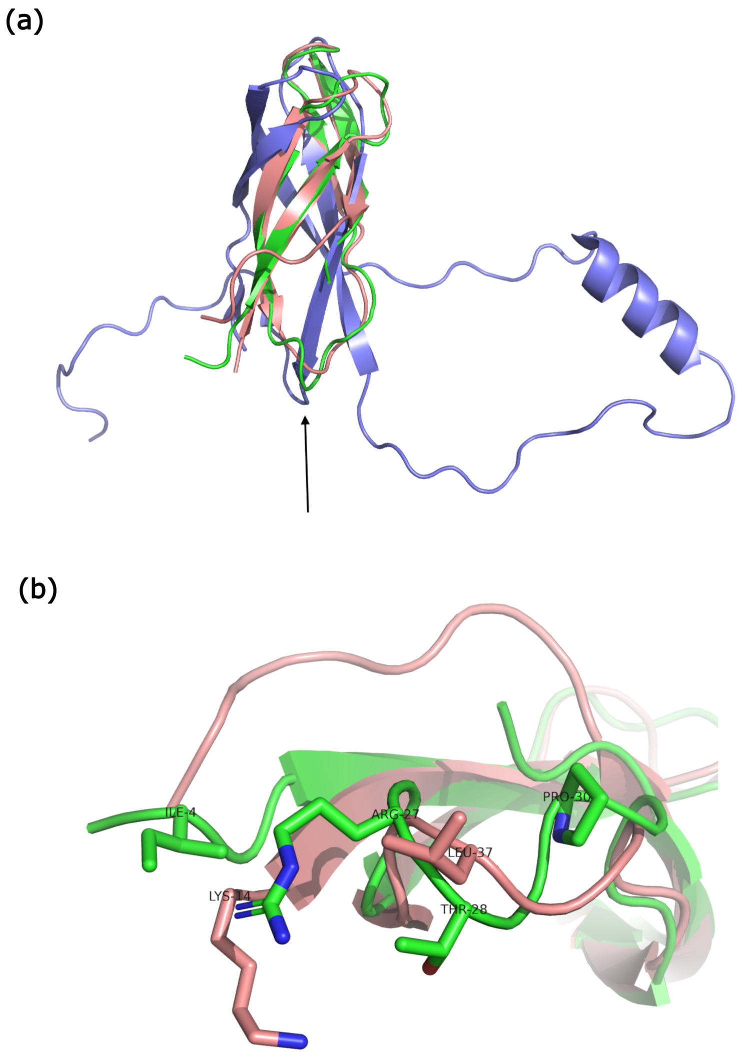


Figure S1: **Bacterial L33 and equivalent proteins in human ribosomes.**
(a) L33 protein from the human mitoribosome (PDB IS 7A5K[1] in mauve), the L36A in the human cytosolic ribosome (PDB IS 8JDM[2] in purple), and E. coli ribosome (PDB IS 4V9D[3] in green) The arrow indicates to the region of interest for this study (around R27 in E. coli). This region has a distinct backbone structure for the human cytosolic ribosome. (b) Zoomed-in perspective of human mitoribosome and E. coli proteins. Residues distinct between the two sequences are labelled, with some E. coli residues lacking human mitoribosome equivalents.

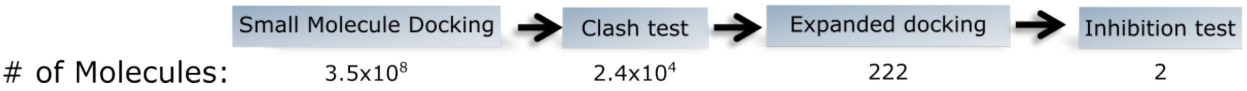


Figure S2: **Summary of the workflow for this study.**
This study consisted of multiple steps, each of which is labelled above with the number of small molecules considered for each step.

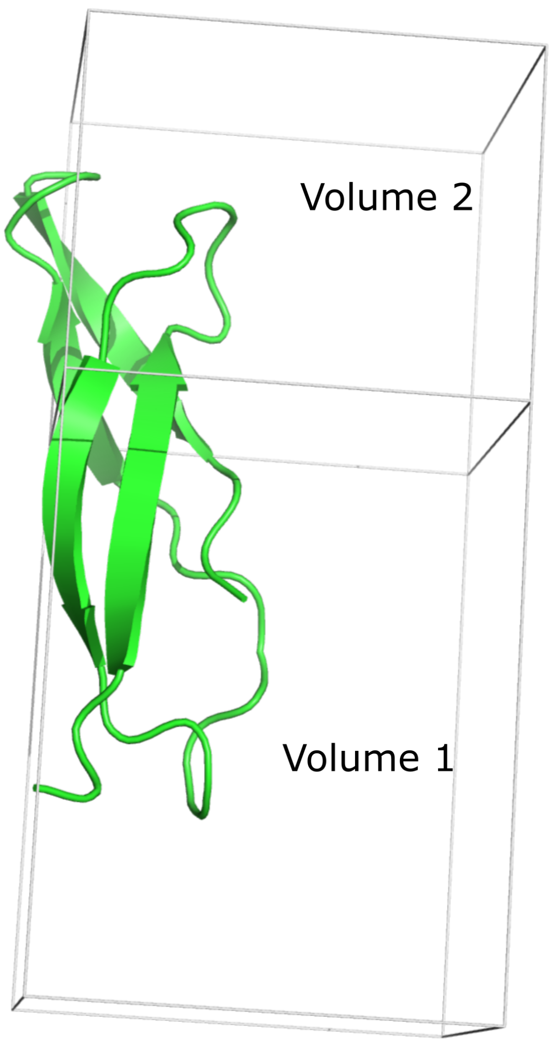


Figure S3: **Boxes used for molecule docking in this work.**
Volume 1 was used for the first round of molecular docking. Both volumes were used the second round of docking, which was performed for residues that had a sufficiently high clash score.

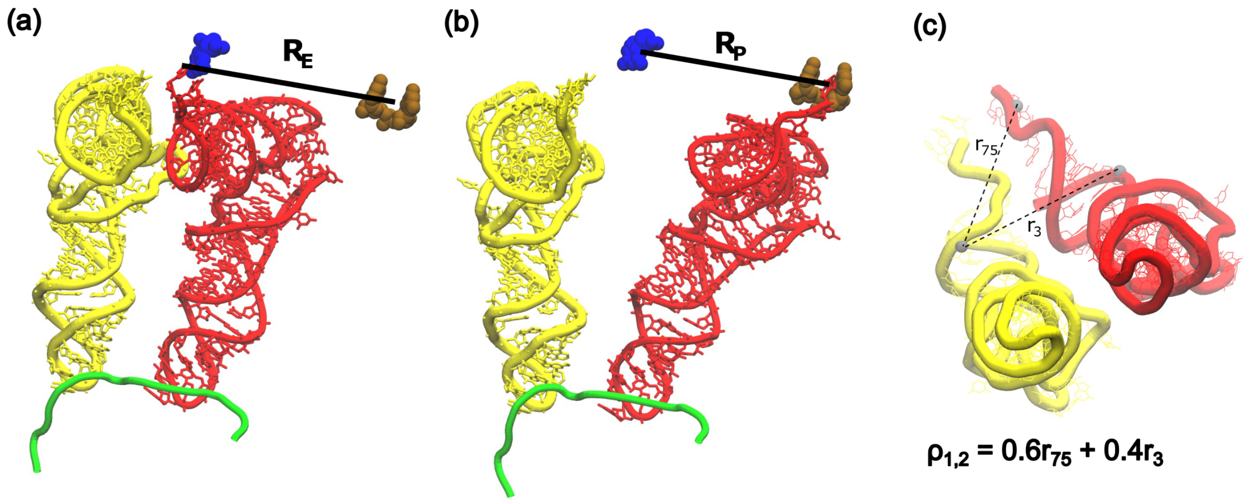


Figure S4: **Reaction coordinates used to detect P/E hybrid formation.**
The coordinates used were previously applied to describe dynamics of P/E formation in a T. Thermophilus ribosome[4]. (a) Representation of the $R_{\mathrm{E}}$ coordinate, the distance between the geometric centers of the C76 nucleobase in the P-site tRNA and the nucleobases of G2421 and A2422 (ochre) on the LSU rRNA. The tRNA is shown in the P/P conformation. (b) Representation of the $R_{\mathrm{P}}$ coordinate, the distance between the geometric centers of the C75 nucleobase in the P-site tRNA and the functional group of G2251 on the rRNA (blue). The tRNA is shown in the P/E conformation. (c) Representation of the $\rho_{1,2}$ coordinate, which is a weighted sum of two interatomic distances: $0.4r_3$+$0.6r_{75}$, where $r_i$ is the distance between O3' atoms of residue i in the P-site tRNA and C67 in the A-site tRNA. The tRNA as shown is in the P/P conformation.

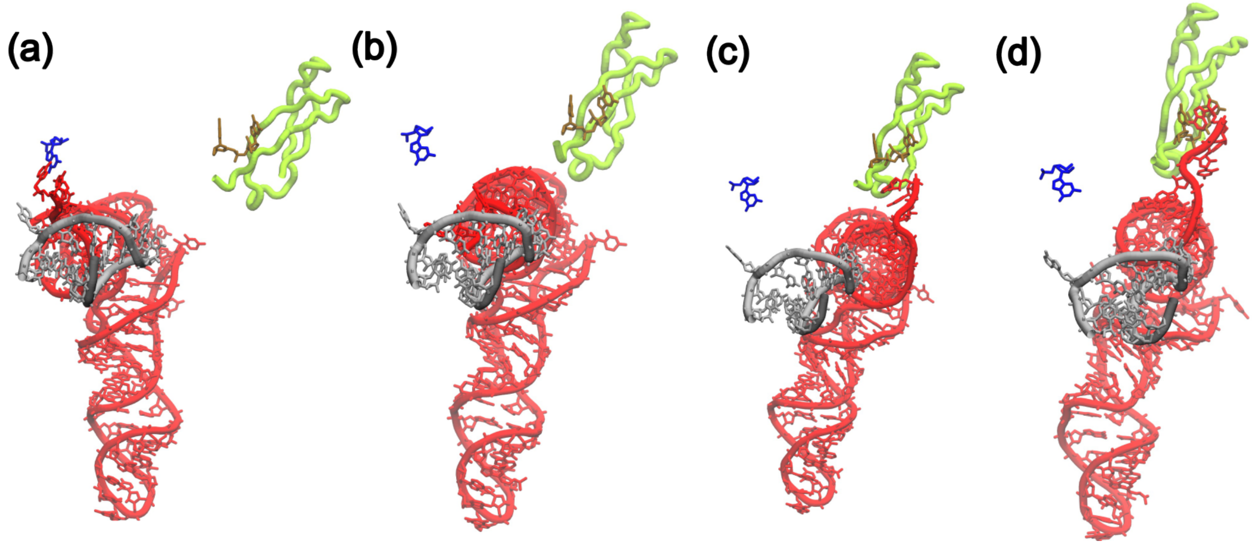


Figure S5: **Intermediates seen along the route to P/E hybrid formation.**
(a) P/P, where the tRNA is firmly in the P site with both elbow and tail in the P site. (b) I1, where the CCA tail has dissociated from the P site but the tRNA has not yet passed H74/H88 of the rRNA (grey) and L33 (green) (c) I2, where the tRNA elbow has passed H74/H88 and L33, but the CCA tail has not yet associated with the E-site residues G2421 and A2422. (d) P/E, where the tRNA elbow and tail have both reached the E site.

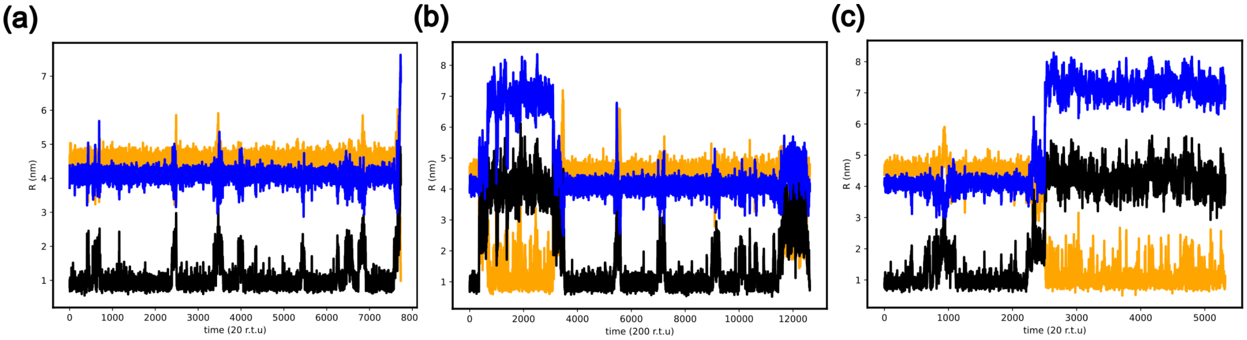


Figure S6: **Representative time traces of SBM trajectories.**
Each panel is a representative run of an SBM with a certain ligand condition. Each reaction coordinate used to monitor P/E hybrid formation is shown ($R_{\mathrm{P}}$, black; $R_{\mathrm{E}}$, orange; $\rho_{1,2}$, blue). a) Simulations with Acl bound. b) Simulations with PPTP bound. c) Simulations of the apo system.

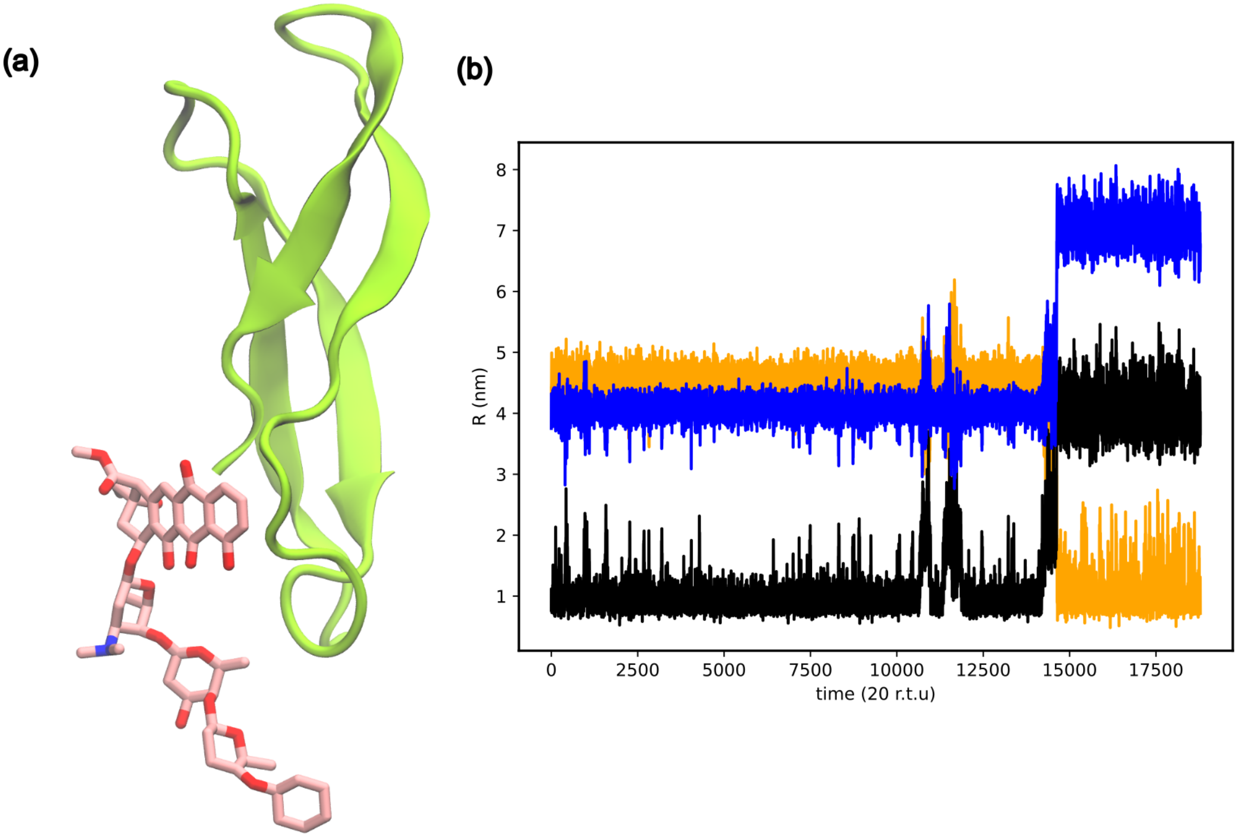


Figure S7: **Acl with a cyclohexyl group attached.**
To demonstrate that addition of a bulk group can further increases the mean first time for P/E hybrid formation, we artificially added a cyclohexyl group to Acl (panel a). (b) Representative time trace of simulations with this ligand, the mean first passage time for P/E hybrid formation is increased by 30%, relative to unmodified Acl.
